# The Mechanism of Neurite Outgrowth Induction by Novel Synthetic Retinobenzoic Acids

**DOI:** 10.1101/2021.03.01.429320

**Authors:** Yang Zhang, Yoji Yoshimi, Osamu Funatsu, Ryuto Hayashi, Shinsuke Komagawa, Shinichi Saito, Yukitoshi Nagahara, Masahiko Ikekita

## Abstract

Retinoids are a family of vitamin A-derived molecules and include the biologically active metabolite, retinoic acid (RA). RA acts as a specific modulator of neuronal differentiation and proliferation. However, teratogenicity and a large excess of RA have been found in animal studies. Thus, development of effective and stable retinoids is desirable. In this study, we showed that treatment with novel synthetic retinobenzoic acids promotes neurite outgrowth in a selected subpopulation of the human neuroblastoma cell line SK-N-SH. Furthermore, we found that, although acting via a different mechanism, retinobenzoic acids have the same neurite outgrowth-inducing effect as RA. Retinoids, including RA, bind to nuclear retinoic acid receptors (RARs). Therefore, we examined the expression of RARs in retinobenzoic acid-treated cells. Similar to already known retinoids, novel synthetic retinobenzoic acids promote the upregulation of RARβ and have no effect on RARα or γ. These results suggest that retinobenzoic acids act via RARβ during neurite outgrowth. Moreover, stimulation with RA or retinobenzoic acids significantly increased the phosphorylation levels of both ERK1/2 and mTOR. ERK1/2 and mTOR inhibition blocked the retinobenzoic acid-induced increase in neurite outgrowth, suggesting that retinobenzoic acids promoted neurite outgrowth by activating the ERK1/2 and mTOR signaling pathways. Notably, the RA-induced increase in neurite outgrowth was blocked by the ERK1/2 inhibitor U0126, but not by the mTOR inhibitor rapamycin. In addition, ERK1/2 inhibition blocked the upregulation of RARβ promoted by RA and retinobenzoic acids. In contrast, mTOR inhibition had no effect on upregulation of RARβ. Our results show that novel synthetic retinobenzoic acids induce neurite outgrowth by a different mechanism than RA. These findings suggest that activation of both ERK1/2, which results in downstream regulation of RARβ, and mTOR, are responsible for the novel synthetic retinobenzoic acid-induced neurite outgrowth in human neuroblastoma cells.

## Introduction

Retinoids are a family of molecules derived from vitamin A and include the biologically active metabolite, retinoic acid (RA). RA regulates multiple biological processes and plays key roles in cell development, growth, and differentiation [1, 2, 3]. For example, RA is known to be crucial in the development of the nervous system [4] and is effective in treatment of many tumors such as acute promyelocytic leukemia [5]. So far, a number of compounds showing retinoidal activity have been reported. However, RA and these traditional synthetic retinoids, containing a hydrocarbon carboxylic acid, exhibit high lipid solubility. Therefore, undesirable side effects like long-lasting teratogenicity and high toxicity are the major drawbacks of retinoid therapy [3, 6]. In an attempt to circumvent this problem, a series of new compounds have been found. Among them, various benzoic acid derivatives elicited specific biological responses similar to those induced by RA, and were thus named retinobenzoic acids [3, 7]. Furthermore, in a pilot study, we found that retinobenzoic acids with a carboxylic acid in the *para* position of the aminobenzoic acid part induce neurite outgrowth.

Neurogenesis plays important roles in numerous processes of the brain, such as learning and memory [8]. Knowing the mechanisms of neurogenesis, including neurite outgrowth, is important for studying brain development and advancing treatments for various neurodegenerative disorders [9]. Neurite outgrowth is necessary for the formation of a functional neuronal network during development. It is also crucial for neuronal plasticity and regeneration [10, 11]. Signaling mechanisms controlling neurite outgrowth have not been well studied and are likely complex processes [11]. Therefore, molecules that promote neurite outgrowth may improve brain development, maintain its function, and have therapeutic potential in the treatment of a wide variety of disorders of the human nervous system.

A large body of evidence supports a role for RA in inducing neurite outgrowth and neuronal differentiation in cultured primary neurons and other model neuronal cell systems, such as dorsal root ganglia [12] or neuroblastoma cells [11, 13, 14]. Therefore, in the present study, we synthesized several novel retinobenzoic acids to examine whether these new compounds have the same effect of inducing neurite outgrowth as RA.

The retinoid signaling pathway has also been shown to be involved in neurite outgrowth [12]. However, the signaling mechanisms of RA that induce neurite outgrowth are only partly known. RA exerts its effects on gene transcription by binding to two families of receptors, the retinoic acid receptors (RARs; α, β, and γ) and the retinoid X receptors (RXRs; α, β, and γ). These receptors belong to the superfamily of nuclear hormone receptors. They are activated by RA, and function as ligand-dependent heterodimeric RAR/RXR transcription factors. RAR/RXR heterodimers control the expression of target genes through binding to specific RA response element (RARE) in promoter regions [3, 11, 15]. The RAR family is activated by all-trans RA (ATRA) and 9-cis retinoic acid (9-cis RA), whereas the RXR family is activated by 9-cis RA [11, 15]. Furthermore, RXR selective ligands alone have no effect on neurite outgrowth in neuroblastoma cells [16].

Among the many different actions of RA in neuroblastoma cells, activation of a number of kinases, including modulation of mitogen-activated protein kinase (MAPK) and phosphoinositide-3-kinase/protein kinase B (PI3K/PKB), has been proposed to play a key role in modulating neurite outgrowth [10, 11]. MAPKs are a family of serine/threonine kinases that mediate signals important for generation of various biological responses. The best-characterized members of the MAPK family are the extracellular signal-regulated kinases, ERK1 and ERK2 [17]. ERKs play an important role in regulating not only neuronal differentiation and survival, but also synaptic plasticity, memory, and learning [11, 18]. Downstream of PI3K, several substrates have been identified that are also likely to play key roles in neurite outgrowth, such as mammalian target of rapamycin (mTOR) [10]. mTOR is a serine/threonine protein kinase and is activated by phosphorylation in response to growth factors, nutrients, mitogens, and hormones [19]. PI3K/Akt/mTOR signaling has been shown to promote neurite growth in cortical neurons [20]. Additional work is needed to understand the relationship between these events and RA/RAR signaling.

Neuroblastoma is a childhood tumor derived from embryonic neural crest cells. Several agents, including RA, induce neuroblastoma cells to differentiate *in vitro*. Therefore, neuroblastoma cells could be a useful model system to study the initial phases of neuronal differentiation [22]. In the present study, we examined the effects of several novel synthetic retinobenzoic acids and investigated their intracellular mechanism of action in a selected neuroblast-like subpopulation, SK-N-SH-N cells [21], derived from the SK-N-SH human neuroblastoma cell line. This manuscript is the first to report that these novel synthetic retinobenzoic acids could promote neurite outgrowth and that this process is mediated by the activation of RARβ, ERKs, and mTOR. More importantly, although retinobenzoic acids are derivatives of and showed the same effect as RA, they act through a different mechanism when inducing neurite outgrowth as compared with RA. In addition, although retinobenzoic acids share similar structures, these compounds showed various effects that were mediated via different mechanisms.

## Materials and methods

### Compounds

Compounds were purchased from the following commercial sources: all-trans retinoic acid (ATRA, Wako Pure Chemicals); 9-cis retinoic acid (9-cis RA, Wako Pure Chemicals); AM580 (Wako Pure Chemicals); LE135 (Tocris Bioscience); U0126 (Cell Signaling Technology); Rapamycin (Sigma-Aldrich). The following retinobenzoic acids were synthesized: SK-78, RH2-36-2, RH2-41-2, RH1-91-2, RH1-95-2, and RH2-22-2. All compounds were dissolved in dimethyl sulfoxide (DMSO). The final concentration (0.1%) of DMSO did not affect the cell viability in SK-N-SH-N cells.

### Cell culture

A selected subpopulation, SK-N-SH-N cells, derived from the SK-N-SH human neuroblastoma cell line [21], were maintained in MEMα medium (Wako Pure Chemicals) supplemented with 10% fetal bovine serum (FBS, Gibco), penicillin (50 U/ml), and streptomycin (0.05 mg/ml) in a 5% CO2 humidified incubator at 37°C.

### Neurite outgrowth analysis

SK-N-SH-N cells were plated onto 6-well tissue culture plates coated with poly-L-lysine (Sigma-Aldrich). Cells were plated at 3×10^5^ cells/ml in MEMα medium containing a very low-serum level (2% FBS), 50 U/ml penicillin and 0.05 mg/ml streptomycin.

Twenty four hours after plating, cells were subjected to stimulation with either one of two retinoic acids or one of seven retinobenzoic acids (6 μM for each compound). For the experiments using inhibitors for RARβ (LE135, 1 or 10 μM), MEK (U0126, 10 μM), and mTOR (Rapamycin, 0.1 μM) SK-N-SH-N cells were pre-treated with these inhibitors 30 min or 24 h prior to ATRA, AM580, RH2-41-2, RH1-95-2, and RH2-22-2 administration (6 μM).

Four days after incubation with retinoic or retinobenzoic acids with or without inhibitors, morphometric analysis was performed on digitized images of live cells taken under phase-contrast illumination, with an inverted microscope (Zeiss, Primo Vert) linked to a digital camera (Canon). The length of neurites not connected to other cells was measured in 5 random fields in each well, with an average of 30 cells per field at a magnification of 200×. Images were analyzed using the ImageJ software.

### Western blot analysis

RARα, β, and γ expression levels and phosphorylation levels of ERK1/2 and mTOR were examined by western blotting in SK-N-SH-N cells that received the following treatments: (1) cells treated with one of two retinoic acids or one of seven retinobenzoic acids (6 μM) for 24 h; (2) cells treated with ATRA, AM580, RH2-41-2, RH1-95-2, and RH2-22-2 (6 μM) for 24 h, 48 h, and 72 h; (3) cells treated with ATRA, AM580, RH2-41-2, RH1-95-2, and RH2-22-2 (6 μM, 24 h or 48 h), and pre-treated for 30 min or 24 h in the presence or absence of a RARβ antagonist (LE135, 1 or 10 μM), an MEK inhibitor (U0126, 10 μM), or an mTOR inhibitor (Rapamycin, 0.1 μM).

Cells were washed in phosphate-buffered saline (PBS), harvested, and solubilized in lysis buffer containing freshly added protease inhibitors (Nacalai Tesque) and phosphatase inhibitors (Nacalai Tesque), and kept on ice for 15 min. Lysates were clarified by centrifugation at 10, 000 rpm for 15 min at 4°C. Proteins were diluted in 1×SDS-PAGE sample buffer (62.5 mM Tris-HCl (pH=6.8)), 5% Glycerol, 1% SDS, 15% 2-mercaptoethanol, 5% bromophenol blue), and heated for 10 min at 100°C. Equal amounts of protein extracts were separated by 7.5-12% SDS-PAGE and transferred onto a Polyvinylidene difluoride (PVDF) membrane (Milipore Corp). The membranes were blocked with 5% non-fat milk in TBS-T (20 mM Tris-HCl, pH 7.5, 150 mM NaCl, 0.05% Tween 20), Blocking One (Nacalai Tesque) or Blocking One-P (Nacalai Tesque) for 30 min at room temperature. These blocked membranes were incubated overnight at 4°C with antibodies against RARα (Abcam), RARβ (Santa Cruz), RARy (Santa Cruz), phospho-ERK1/2 (Cell Signaling Technology), or phospho-mTOR (Cell Signaling Technology) as primary antibodies. These primary antibodies were diluted in immunoreaction enhancer solution 1 (Can Get Signal, TOYOBO). The membranes were then washed twice with TBS-T followed by incubation for 1 h at room temperature in immunoreaction enhancer solution 2 (Can Get Signal; TOYOBO) with peroxidase-conjugated antibody against rabbit IgG (Dako) as the secondary antibody. The membranes were then washed three times with TBS-T, washed twice with TBS (20 mM Tris-HCl, pH 7.5, 150 mM NaCl), and immunoreactivity was detected by the ECL Prime Western Blotting Detection reagent (GE Healthcare Bioscience). Images were captured and immunoreactive bands were quantified using a ChemiDoc MP ImageLab PC system (Bio-Rad). Membranes were stripped in a 62.5 mM Tris buffer containing 2% SDS and 100 mM β-mercaptoethanol (pH 6.7), and then the membranes were re-probed with anti-ERK1/2 (Cell Signaling Technology), or anti-mTOR (Cell Signaling Technology) using Can Get Signal. GAPDH (Sigma-Aldrich) immunoreactivity was used to monitor equal sample loading. Western blot results presented here are representative of at least three independent experiments.

## Results

### Retinobenzoic acids increase neurite outgrowth in SK-N-SH-N cells

In a past study, RA had the ability to induce neurite outgrowth [11]. Thus, to find out whether novel synthetic retinobenzoic acids (Figure 1) have the same effect of inducing neurite outgrowth as RA, we examined the morphological changes induced by RA and retinobenzoic acids in SK-N-SH-N cells. Neurite outgrowth analysis was performed by determining and analyzing the length of neurites not connected to the other cells. As shown in Figure 2, RA and four retinobenzoic acids AM580, SK-78, RH2-36-2, and RH2-41-2 promoted neurite outgrowth in SK-N-SH-N cells within 96 h, when compared with DMSO-treated control cells. In addition, a novel synthetic retinobenzoic acid RH2-41-2 showed a highly similar effect of inducing neurite outgrowth as RA.

**Figure 1.**
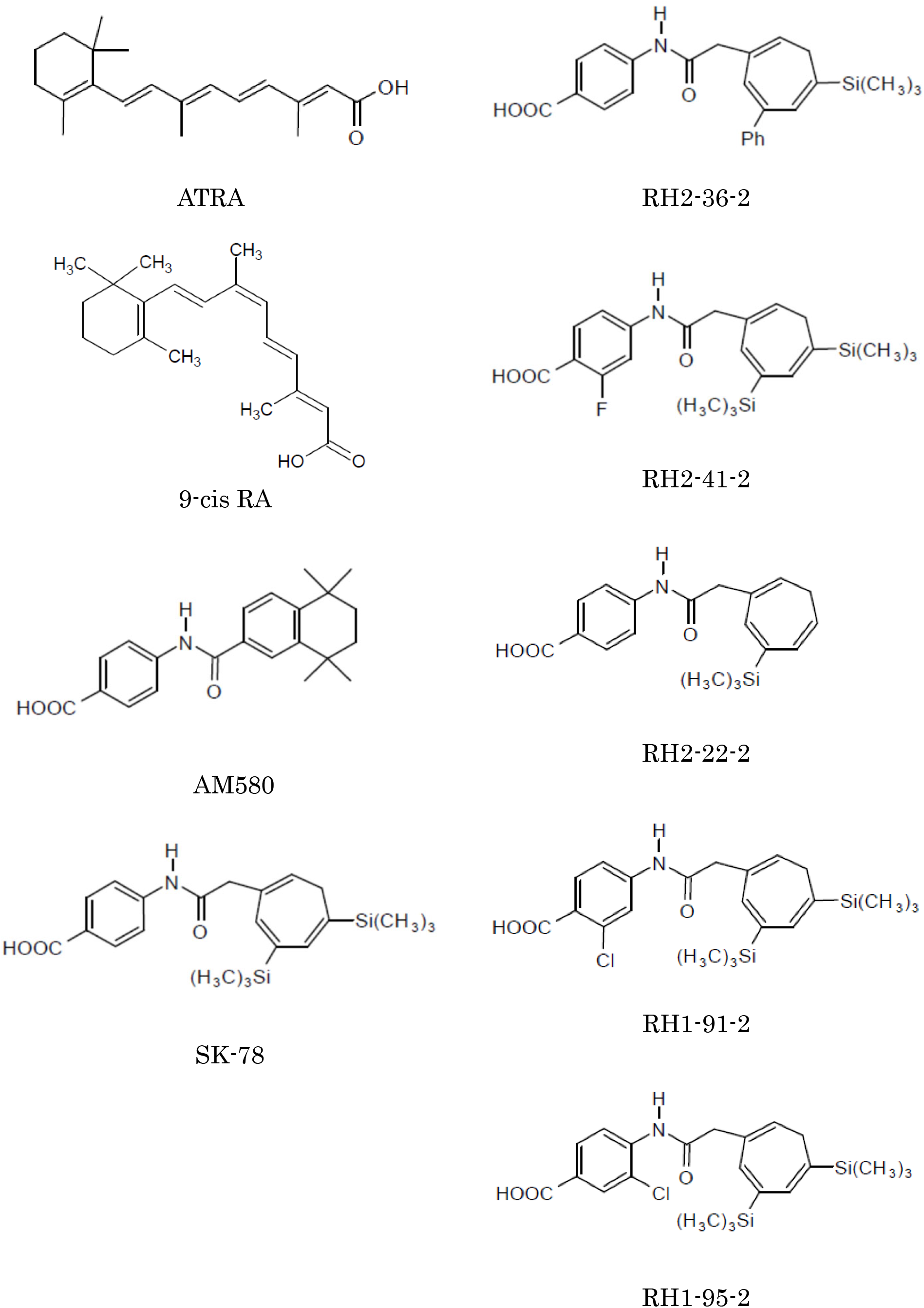
Chemical structures of ATRA, 9-cis RA, and the synthetic retinobenzoic acids.

**Figure 2.**
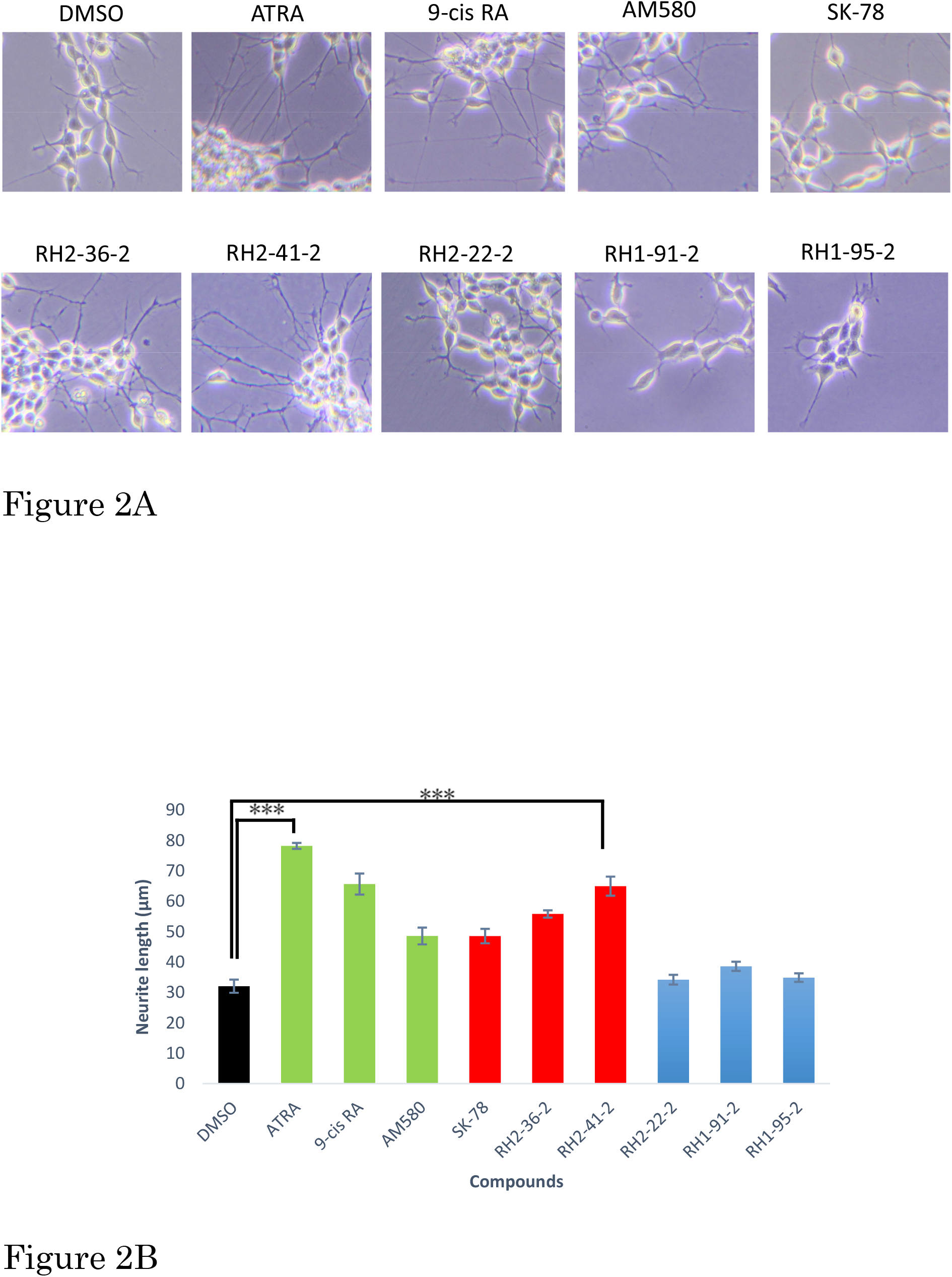
RA and four retinobenzoic acids promote neurite outgrowth in SK-N-SH-N cells. (A) Representative images of SK-N-SH-N cells after indicated treatments. Neurite outgrowth was analyzed after 96 h of incubation with compounds at 6 μM. Cell morphology was observed under a light microscope (200×). (B) Quantification of the measured neurite length. Results are expressed as means ± SEM (n = 7). ***p < 0.001 versus control cells.

### Retinobenzoic acids promote the upregulation of RARβ

Retinoids, including RA, exert their effects on gene transcription by binding to nuclear RARs, and among RA-responsive target genes include RARs themselves [3, 11, 15, 16]. Thus, to determine the neurite outgrowth-inducing mechanism of RA and retinobenzoic acids in SK-N-SH-N cells, the expression level of three subtypes of RARs (α, β, and γ) was examined. As show in Figure 3, treatment with RA or retinobenzoic acids AM580, SK-78, RH2-36-2, RH2-41-2 and RH2-22-2 significantly promoted the upregulation of RARβ and had no effect on RARα or γ at both mRNA (data not shown) and proteins level. These results suggest a role for RARβ in the retinobenzoic acids-induced outgrowth of neurites. Interestingly, though RH2-22-2 had no ability of inducing neurite outgrowth, it significantly promoted the upregulation of RARβ. Therefore, we assumed that the upregulation of RARβ is a necessary condition for the promotion of retinobenzoic acids-induced neurite outgrowth.

**Figure 3.**
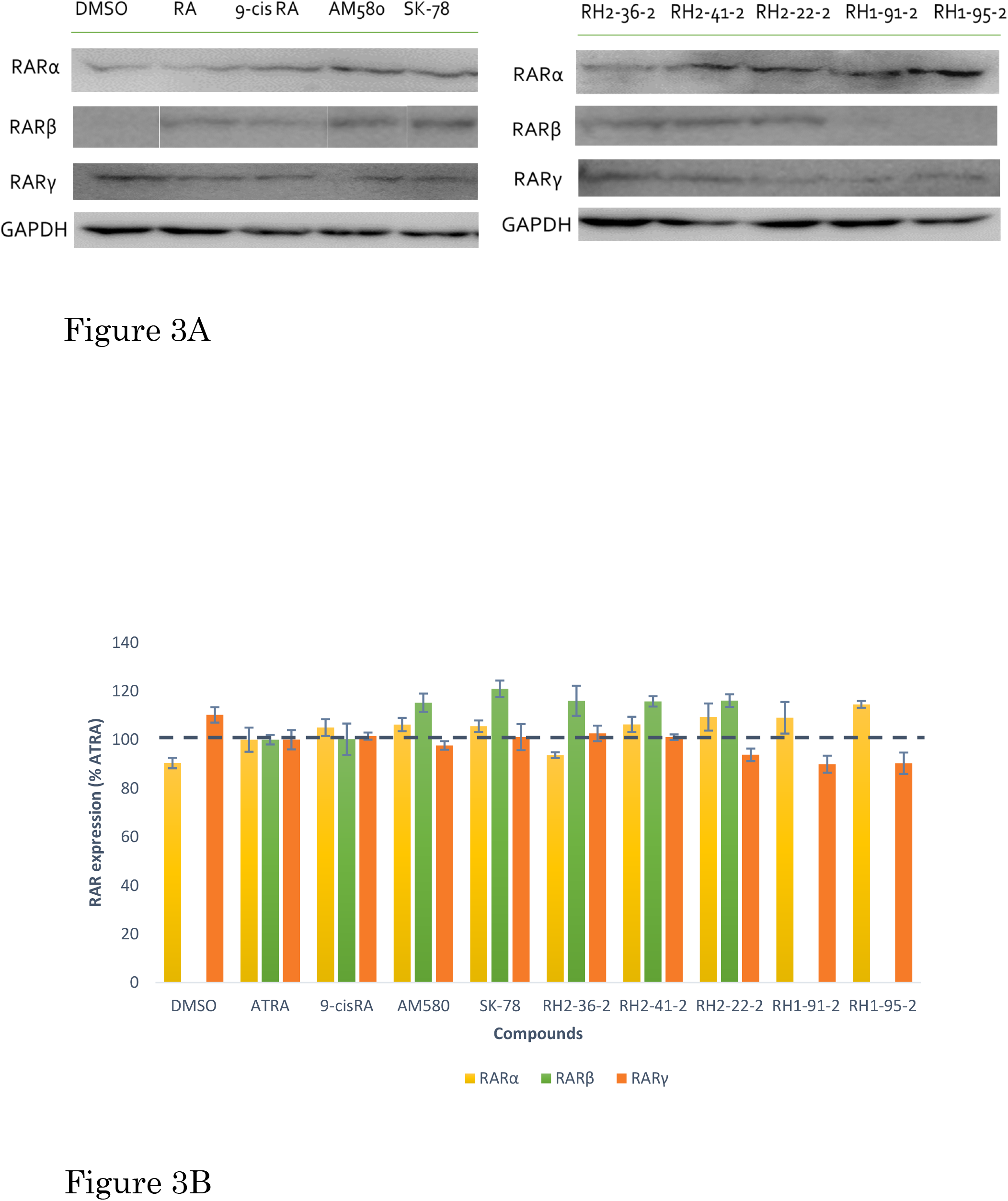
The expression level of retinoic acid receptors (RARs) in RA- and retinobenzoic acid-treated cells. SK-N-SH-N cells were treated with ATRA, 9-cis RA, AM580, SK-78, RH2-36-2, RH2-41-2, RH2-22-2, RH1-91-2 and RH1-95-2 (6 μM) for 24 h. (A) Protein extracts were analyzed by western blotting. (B) Data are calculated as the ratio of RARs/GAPDH and are shown as means ± SEM (n = 3).

### AM580- and RH2-41-2-induced neurite outgrowth requires upregulation of RARβ

Five representative compounds were further selected to study the neurite outgrowth-inducing mechanism of RA and retinobenzoic acids in SK-N-SH-N cells. ATRA was chosen as a positive control. We also selected the already characterized retinobenzoic acid AM580; RH2-41-2, which showed an effect of inducing neurite outgrowth highly similar to RA; RH1-95-2, which had no effect on neurite outgrowth; and RH2-22-2, which had no ability of inducing neurite outgrowth but promoted the upregulation of RARβ.

To determine whether the upregulation of RARβ is involved in RA- or retinobenzoic acids-induced neurite growth, we pre-treated SK-N-SH-N cells with the RARβ antagonist LE135 at 1 μM or 10 μM concentration, 24 h prior to ATRA, AM580, RH2-41-2, RH1-95-2 and RH2-22-2 administration. As shown in Figure 4A, LE135 inhibits ATRA- or retinobenzoic acids-induced RARβ upregulation. Figure 4B shows that the inhibition of RARβ upregulation by LE135 administration significantly reduced AM580- and RH2-41-2-induced neurite outgrowth. However, LE135 treatment only reduced ATRA-induced neurite outgrowth to a small extent that was not significant. These results suggest that RARβ upregulation is involved in AM580 and RH2-41-2-induced neurite outgrowth in SK-N-SH-N cells. Moreover, it is impossible to definitely state that upregulation of RARβ is not involved in ATRA-induced neurite outgrowth. We consider that there is a possibility that the pathway of ATRA-induced neurite outgrowth is complicated, and that inhibition of RARβ upregulation alone is insufficient to abolish ATRA-induced neurite outgrowth in SK-N-SH-N cells.

**Figure 4.**
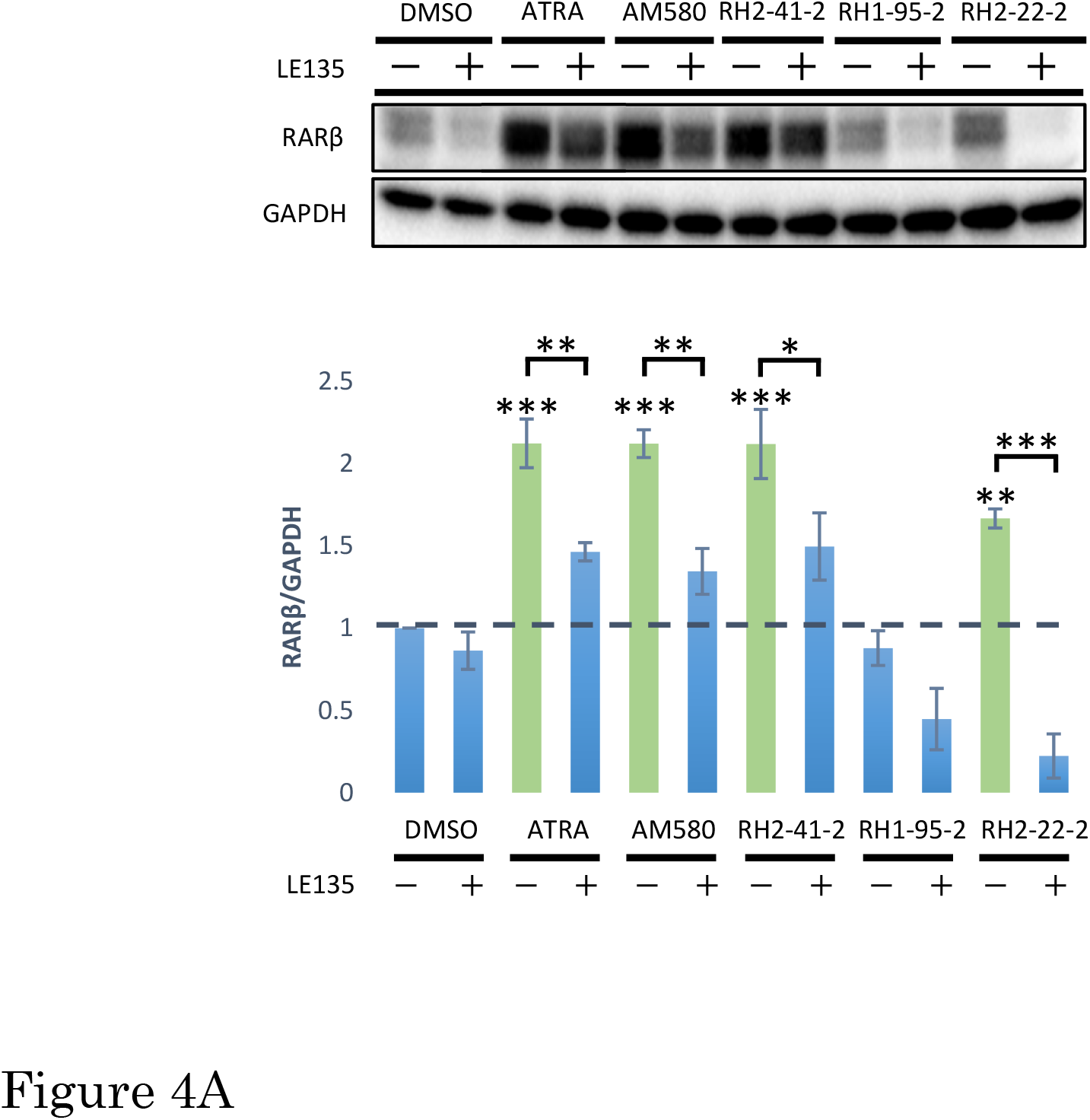
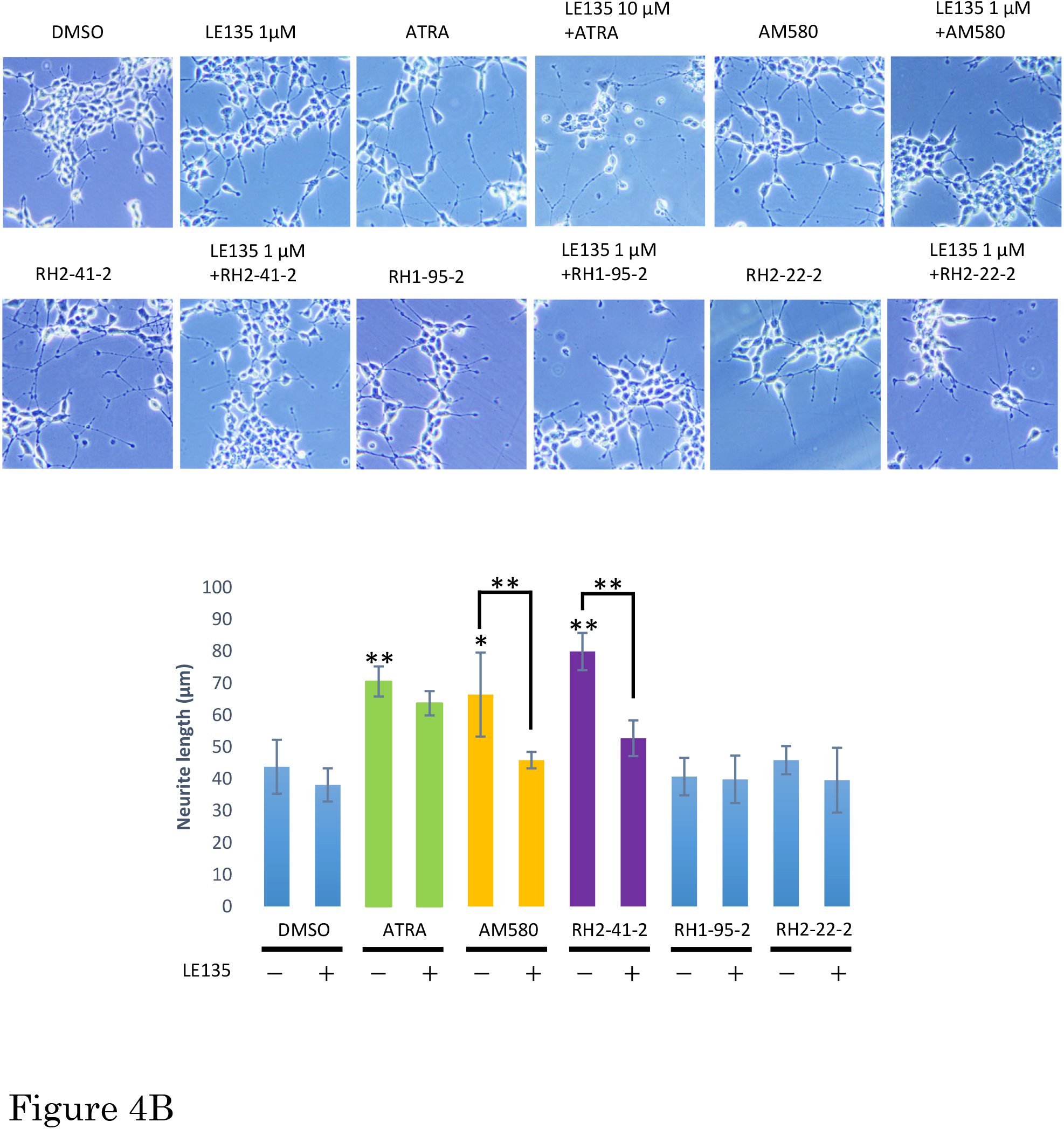
Effect of RARβ antagonist LE135 on ATRA- or retinobenzoic acids-induced upregulation of RARβ and neurite outgrowth. (A) LE135 inhibits ATRA- or retinobenzoic acids-induced RARβ activation. SK-N-SH-N cells were pretreated with 1 μM or 10 μM LE135 24 h prior to ATRA or retinobenzoic acids exposure (6 μM). 24 h following ATRA or retinobenzoic acids stimulation, cellular extracts were prepared and analyzed by western blotting. Quantified protein expression levels are expressed as fold change from DMSO treated cells. *** p < 0.001 compared to control cells or as indicated. (B) LE135 has no effect on ATRA-induced neurite outgrowth, but inhibits retinobenzoic acids-induced neurite outgrowth. SK-N-SH-N cells were pretreated with LE135 24 h prior to ATRA or retinobenzoic acids exposure (6 μM), and 96 h following ATRA or retinobenzoic acids treatment, morphology was examined and neurite length was measured under a light microscope (200×). Results are expressed as means ± SEM (n = 3). ***p < 0.001, **p < 0.01, *p < 0.05 versus control cells.

### AM580- and RH2-41-2-induced neurite outgrowth requires activation of ERK1/2

Extracellular signal-regulated kinases (ERK1/2) are well known to play an essential role in modulating neurite outgrowth [11, 14, 23, 24, 25]. To determine whether ATRA or retinobenzoic acids induce ERK1/2 activation in SK-N-SH-N cells, expression levels of phosphorylated forms of ERK1/2 were investigated by immunoblotting. Furthermore, the role that ERK1/2 pathway activation plays in ATRA-, AM580-, and RH2-41-2-induced neurite outgrowth was examined with the MEK inhibitor U0126. We pre-treated SK-N-SH-N cells with 10 μM U0126 30 min prior to ATRA, AM580, RH2-41-2, RH1-95-2 and RH2-22-2 administration. As shown in Figure 5A, U0126 inhibits ATRA- or retinobenzoic acids-induced ERK1/2 phosphorylation. Figure 5B shows that the inhibition of ERK1/2 phosphorylation by U0126 treatment significantly reduced ATRA-, AM580-, and RH2-41-2-induced neurite outgrowth. These results suggest that ERK1/2 activation is involved in ATRA-, AM580- and RH2-41-2-induced neurite outgrowth in SK-N-SH-N cells.

**Figure 5.**
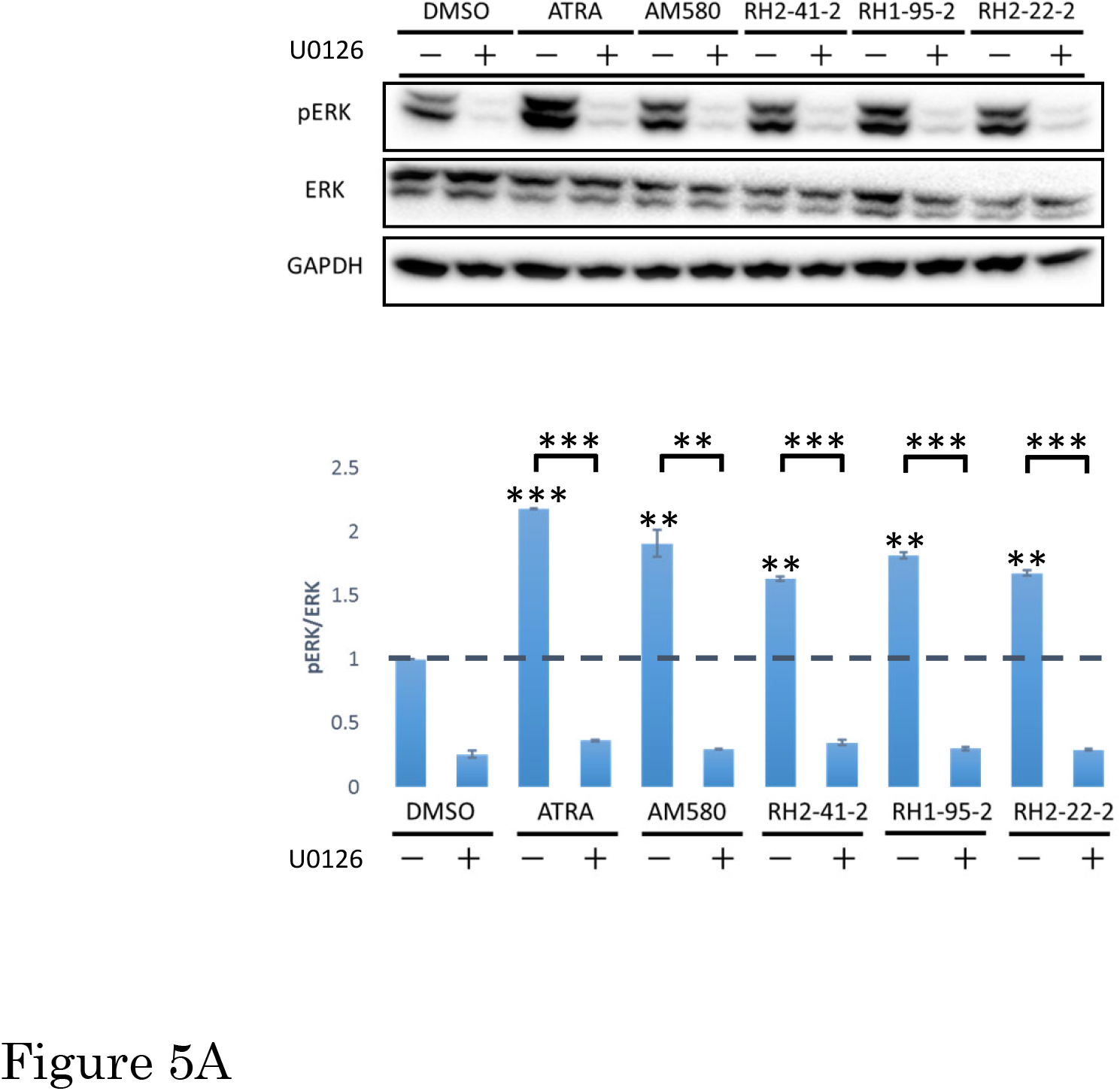
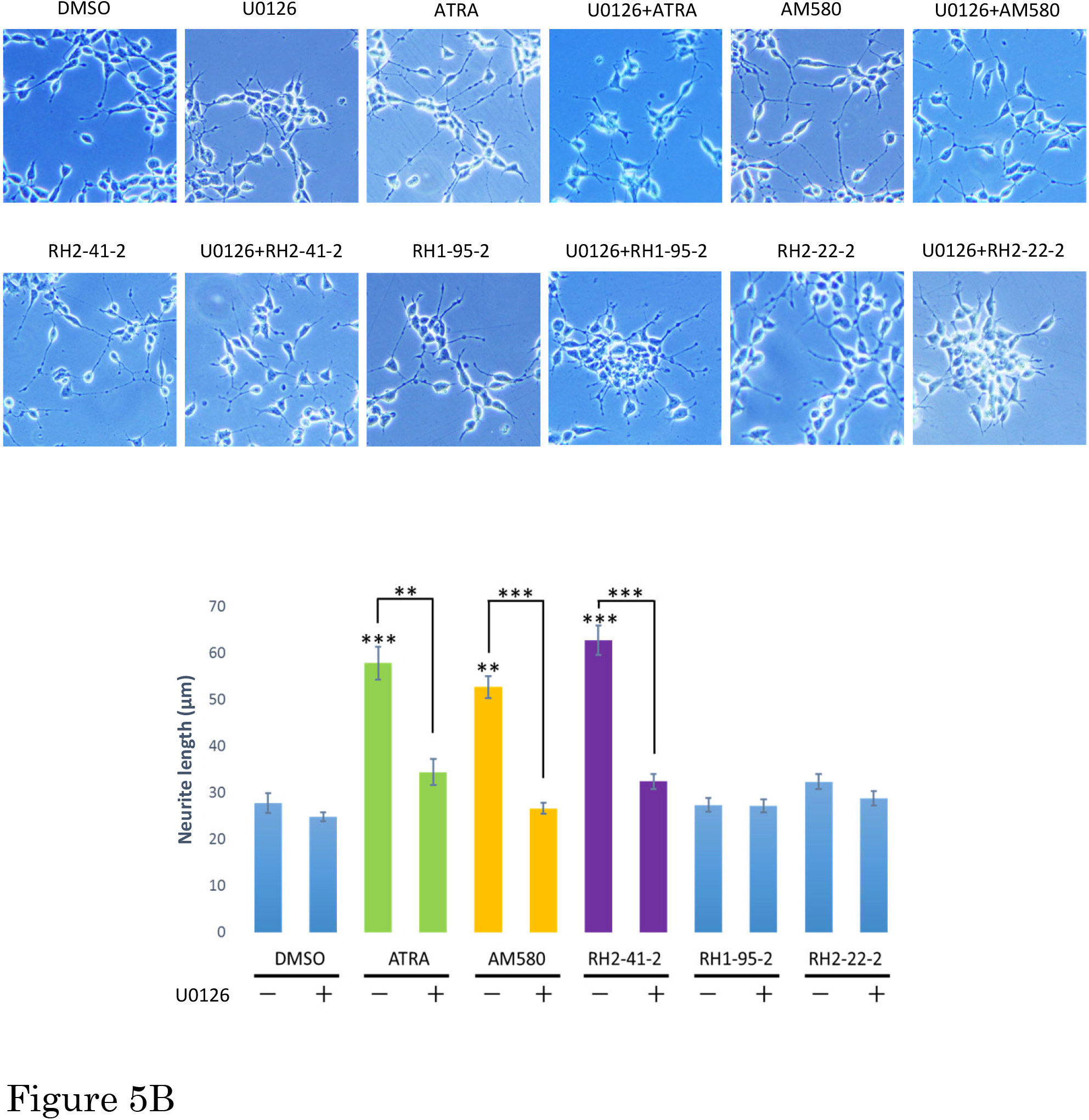
Effect of MEK inhibitor U0126 on ATRA- or retinobenzoic acids-induced ERK1/2 phosphorylation and neurite outgrowth. (A) U0126 inhibits ATRA- or retinobenzoic acids-induced ERK1/2 phosphorylation. SK-N-SH-N cells were pre-treated with 10 μM U0126 30 min prior to ATRA or retinobenzoic acids exposure (6 μM), and 24 h following ATRA or retinobenzoic acids stimulation, cellular extracts were prepared and analyzed by western blotting. Quantified protein expression levels are expressed as fold change from DMSO treated cells. (B) U0126 inhibits ATRA- or retinobenzoic acids-induced neurite outgrowth. SK-N-SH-N cells were pre-treated with 10 μM U0126 30 min prior to ATRA or retinobenzoic acids exposure (6 μM), and 96 h following ATRA or retinobenzoic acids treatment, morphology was examined and neurite length was measured under a light microscope (200×). Results are expressed as means ± SEM (n = 3). ***p < 0.001, **p < 0.01 versus control cells or as indicated.

### Extracellular signal-regulated kinases (ERK1/2) act upstream of RARβ

Both ERK1/2 and RARβ activation is involved in AM580- and RH2-41-2-induced neurite growth in SK-N-SH-N cells. To elucidate the relationship between ERK1/2 and RARβ, the expression level of RARβ in the presence of U0126 was examined. When cells were pre-treated with U0126, ATRA-, AM580-, and RH2-41-2-induced RARβ upregulation was substantially decreased (Figure 6). These results suggest that ERK1/2 acts upstream of RARβ in inducing neurite outgrowth.

**Figure 6.**
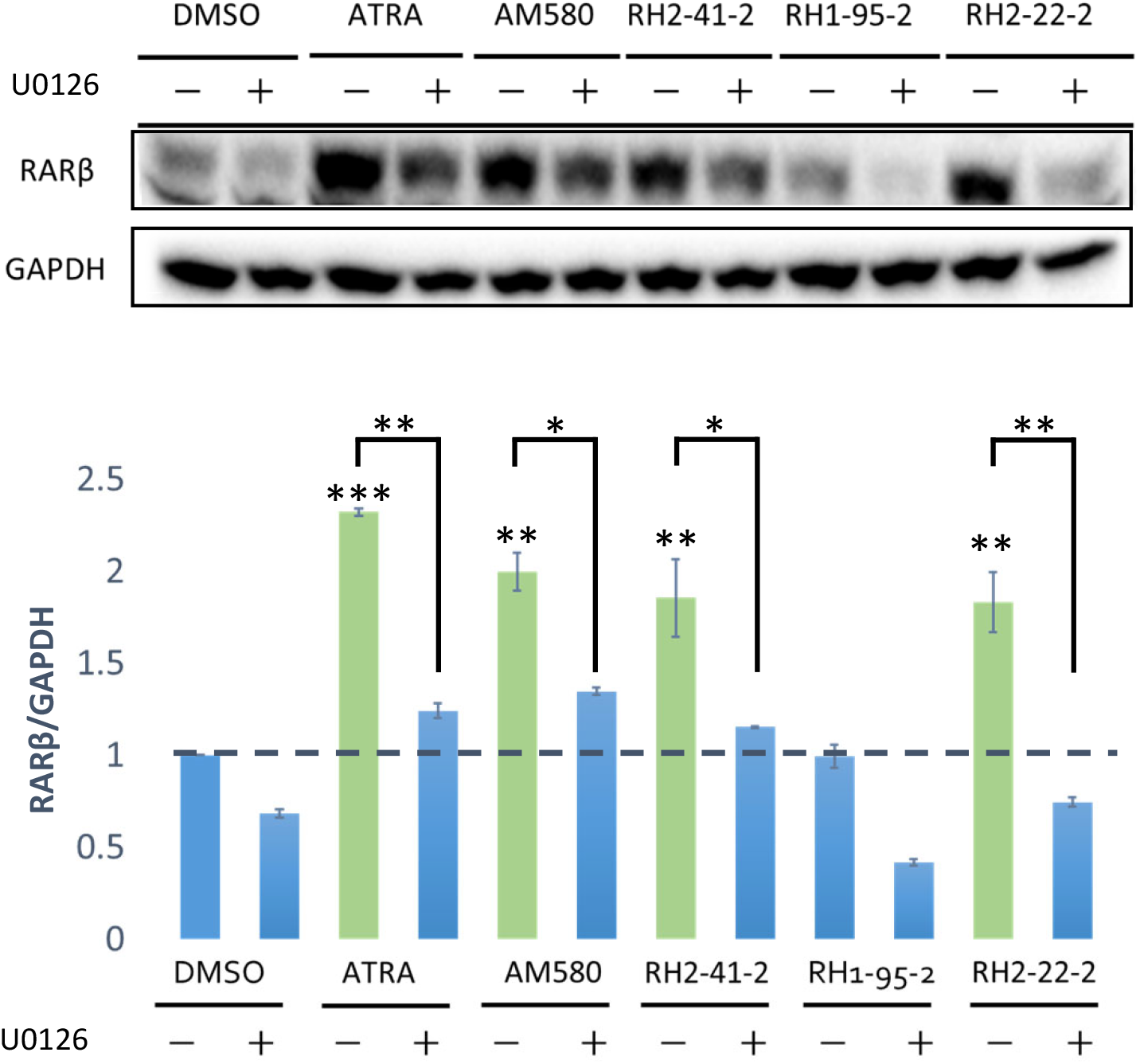
MEK inhibitor U0126 inhibits ATRA- or retinobenzoic acids-induced upregulation of RARβ. SK-N-SH-N cells were pre-treated with 10 μM U0126 30 min prior to ATRA or retinobenzoic acids exposure (6 μM), and 24 h following ATRA or retinobenzoic acids stimulation, cellular extracts were prepared and analyzed by western blotting. Data are calculated as the ratio of RARβ/GAPDH and are shown as means ± SEM (n = 3). ***p < 0.001, **p < 0.01, *p < 0.5 versus control cells or as indicated.

### AM580- and RH2-41-2-induced neurite outgrowth requires activation of mTOR

The cross talk of Ras–extracellular signal-regulated kinase (Ras-ERK) and phosphatidylinositol-3-kinase–mammalian target of rapamycin (PI3K-mTOR) signaling pathways plays an essential role in cell survival, differentiation and proliferation [26]. Furthermore, recent studies have indicated that PI3K/Akt/mTOR signaling is involved in neuronal differentiation [15, 24]; for example, atorvastatin enhances neurite outgrowth via mTOR signaling pathways [20]. To determine whether ATRA or retinobenzoic acids activate mTOR in SK-N-SH-N cells, expression levels of phosphorylated forms of mTOR were examined by immunoblotting. Furthermore, we investigated the role of mTOR pathway activation on ATRA-, AM580-, and RH2-41-2-induced neurite outgrowth by using the mTOR inhibitor rapamycin. SK-N-SH-N cells were treated with 0.5 μM rapamycin 30 min prior to ATRA, AM580, RH2-41-2, RH1-95-2, and RH2-22-2 administration. As shown in Figure 7A, rapamycin inhibited ATRA- or retinobenzoic acids-induced mTOR phosphorylation. Figure 7B shows that the inhibition of mTOR phosphorylation by rapamycin treatment significantly reduced AM580- and RH2-41-2-induced neurite outgrowth. However, ATRA-induced neurite outgrowth was only reduced by rapamycin to a small extent that was not significant. These results suggest that mTOR activation is involved in AM580 and RH2-41-2-induced neurite outgrowth in SK-N-SH-N cells. Conversely, there is a need for further studies investigating whether ATRA-induced mTOR phosphorylation is related to ATRA-induced neurite outgrowth.

**Figure 7.**
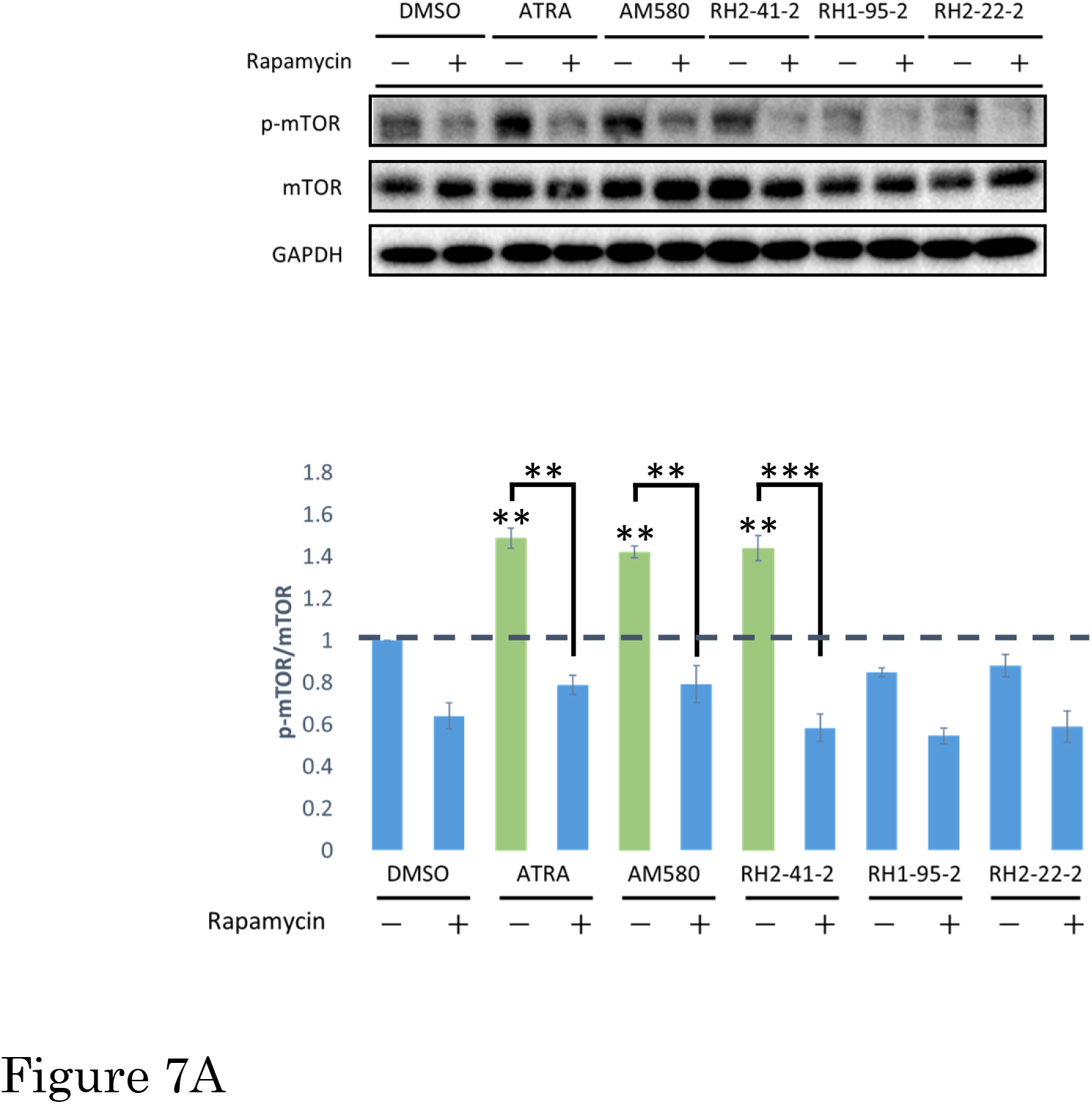
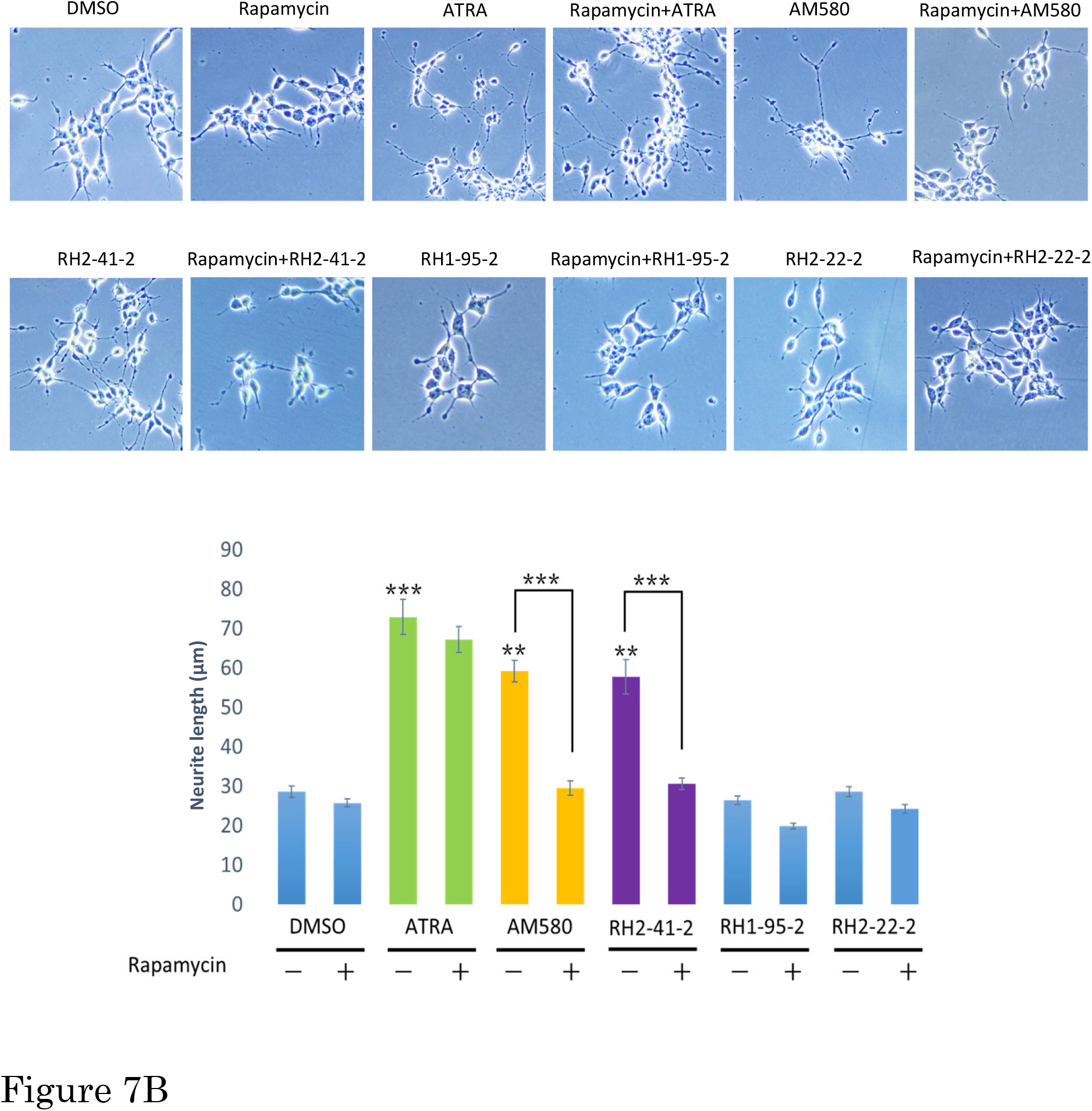
Effect of mTOR inhibitor rapamycin on ATRA- or retinobenzoic acids-induced mTOR phosphorylation and neurite outgrowth. (A) Rapamycin inhibits ATRA- or retinobenzoic acids-induced mTOR phosphorylation. SK-N-SH-N cells were pretreated with 0.5 μM rapamycin 30 min prior to ATRA or retinobenzoic acids exposure (6 μM), and 24 h following ATRA or retinobenzoic acids stimulation, cellular extracts were prepared and analyzed by western blotting. Quantified protein expression levels are expressed as fold change from DMSO treated cells. (B) Rapamycin has no effect on ATRA-induced neurite outgrowth, but inhibits retinobenzoic acids-induced neurite outgrowth. SK-N-SH-N cells were pretreated with 0.5 μM rapamycin 30 min prior to ATRA or retinobenzoic acids exposure (6 μM), and 96 h following ATRA or retinobenzoic acids treatment, morphology was examined and neurite length was measured under a light microscope (200×). Results are expressed as means ± SEM (n = 3). ***p < 0.001, **p < 0.01 versus control cells or as indicated.

### Mammalian target of rapamycin (mTOR) does not act upstream of RARβ

We next sought to determine whether mTOR acts upstream of RARβ. To this end, the expression levels of RARβ in the presence of rapamycin were examined. When cells were pre-treated with rapamycin, rapamycin did not affect ATRA- or retinobenzoic acids-induced upregulation of RARβ (Figure 8). These results suggest that mTOR does not act upstream of RARβ in inducing neurite outgrowth.

**Figure 8.**
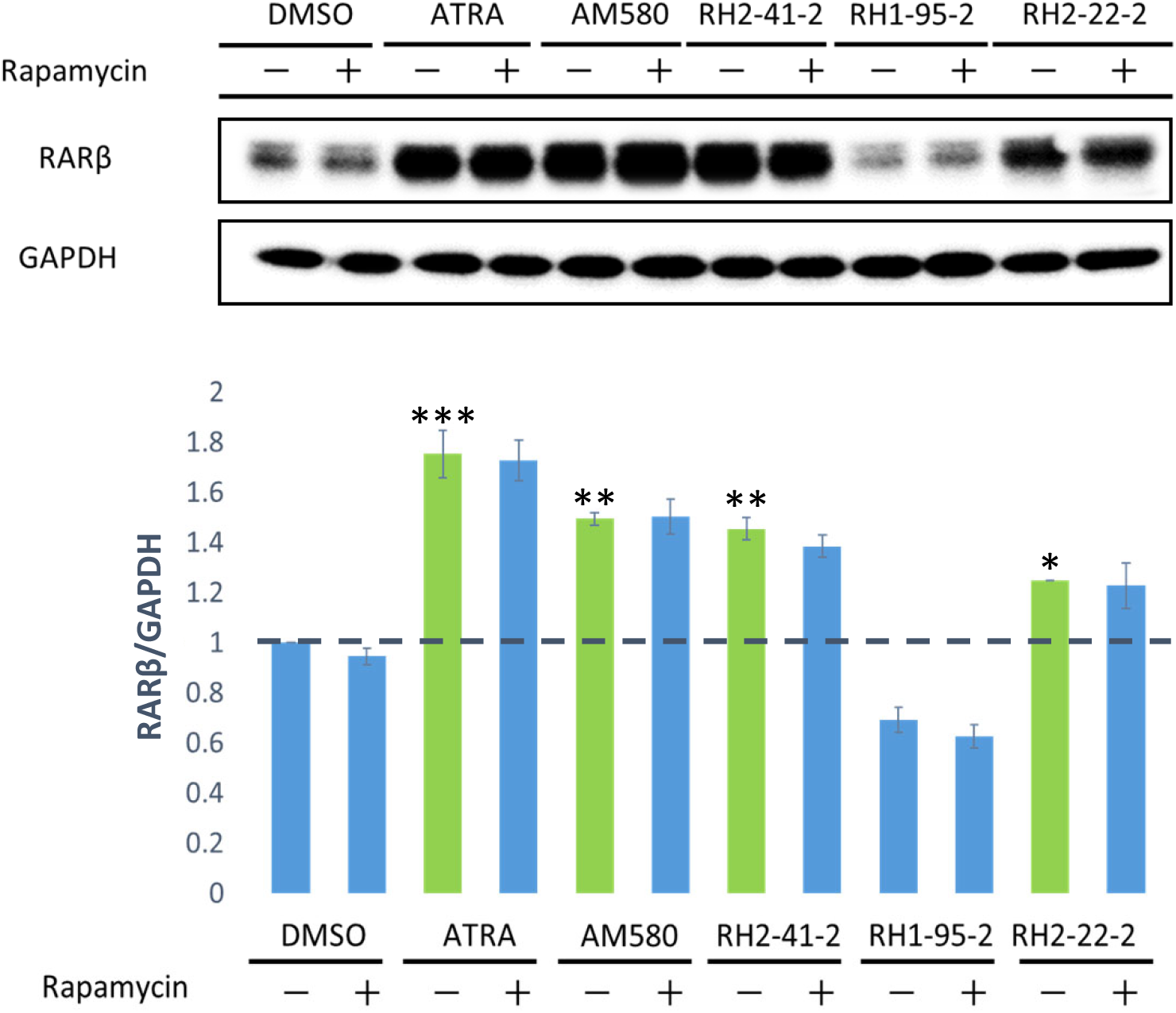
mTOR inhibitor rapamycin has no effect on ATRA or retinobenzoic acids-induced upregulation of RARβ. SK-N-SH-N cells were pretreated with 0.5 μM rapamycin 30 min prior to ATRA or retinobenzoic acids exposure (6 μM), and 24 h following ATRA or retinobenzoic acids stimulation, cellular extracts were prepared and analyzed by western blotting. Data are calculated as the ratio of RARβ/GAPDH and are shown as means ± SEM (n = 3). ***p < 0.001, **p < 0.01, *p < 0.05 versus control cells.

## Discussion

In this study, we demonstrated that treatment with novel synthetic retinobenzoic acids could promote neurite outgrowth in SK-N-SH-N cells. Retinobenzoic acids involve a different mechanism of inducing neurite outgrowth as compared with RA. In addition, although retinobenzoic acids have similar structures, these compounds showed various effects mediated by different mechanisms.

This report is the first to demonstrate that these novel synthetic retinobenzoic acids could promote neurite outgrowth by activating both ERK1/2, which results in downstream upregulation of RARβ and mTOR. In contrast to the mechanism of retinobenzoic acids, ATRA induced neurite outgrowth by activation of ERK1/2, but not mTOR.

RARβ activation has been reported to play a role in mediating neuronal differentiation, including neurite outgrowth [11, 16, 27]. In the present study, ATRA stimulation activated the RARβ signaling pathway. However, treatment with LE135 that inhibits RARβ activity did not abolish ATRA-induced neurite outgrowth. It is impossible to conclude that upregulation of RARβ is not involved in ATRA-induced neurite outgrowth. We consider that there is a possibility that the pathway of ATRA-induced neurite outgrowth is complicated, and that the inhibition of RARβ upregulation alone is insufficient to abolish ATRA-induced neurite outgrowth in SK-N-SH-N cells. In addition, RA can induce apoptosis through caspase activation via pathways independent of the nuclear receptor, in which retinoid-related molecules do not bind to classical retinoid receptors [28]. It could thus be possible that ATRA-induced neurite outgrowth was mediated by pathways independent from nuclear receptors. Stimulation with the traditional synthetic retinobenzoic acid AM580 and the novel synthetic retinobenzoic acid RH2-41-2 activated RARβ, and inhibition of RARβ activity abolished AM580- and RH2-41-2-induced neurite outgrowth. These data indicate that RARβ activation is involved in AM580- and RH2-41-2-induced neurite outgrowth in SK-N-SH-N cells. Notably, although novel synthetic retinobenzoic acid RH2-22-2 was unable to induce neurite outgrowth, it significantly promoted the upregulation of RARβ, suggesting that RARβ upregulation is a necessary condition for the promotion of retinobenzoic acids-induced neurite outgrowth. Furthermore, in contrast to ATRA, which only binds to RARs, AM580 can bind to RARs and RXRs. Thus, it is possible that RH2-22-2 was unable to induce neurite outgrowth similarly to other retinobenzoic acids, as it could not bind to RXRs.

Administration of the MEK inhibitor U0126, which could inhibit ERK1/2 phosphorylation, significantly prevented ATRA-, AM580-, and RH2-41-2-induced neurite outgrowth. These results provide strong evidence that the activation of ERK1/2 is required for ATRA-, AM580-, or RH2-41-2-induced neurite outgrowth in SK-N-SH-N cells. In addition, ERK1/2 is well known to be involved in modulating neurite outgrowth. For example, ERK1/2 activation played an important role in neurite outgrowth induced by α-lipoic acid and honokiol [23, 25]. Consequently, our data suggest that ERK1/2 activation is essential for neurite outgrowth. However, the mechanism by which ATRA, AM580, or RH2-41-2 activates the ERK1/2 signaling pathway is still unclear. Furthermore, additional work is needed to understand the relationship between ERK1/2 activation and RARβ signaling. After the inhibition of ERK1/2, ATRA-, AM580-, or RH2-41-2-induced RARβ upregulation was decreased. These results suggest that ERK1/2 acts upstream of RARβ in inducing neurite outgrowth. The relationship between kinase activation and RAR signaling is actually not well understood. A recent study revealed that RAR activation could lead to MAPK/ERKs phosphorylation. After nuclear translocation these enzymes phosphorylate nuclear cofactors and bind to RAR/RXR heterodimers, converging on MSK-1, of histone 3 (H3), eventually affecting the regulation of gene transcription [29]. In consequence, we consider that there could be a feedback route between RARβ and ERK1/2, which should be analyzed in the future.

Several substrates have been identified downstream of PI3K that are likely to play key roles in neurite outgrowth, such as mTOR [10]. Moreover, recent studies have indicated that PI3K/Akt/mTOR signaling is involved in neuronal differentiation [15, 24]. In the present study, ATRA, AM580 or RH2-41-2 treatment activated mTOR signaling. However, inhibition of mTOR activation by rapamycin abolished AM580- and RH2-42-3-, but not ATRA-stimulated neurite outgrowth. These results suggest that mTOR activation is required for AM580- or RH2-41-2-induced neurite outgrowth in SK-N-SH-N cells. Rapamycin treatment reduced ATRA-induced neurite outgrowth to a small extent that was not significant. Taking into account the fact that rapamycin is an inhibitor of mTORC1; these results could suggest different roles of mTOR in promoting neurite outgrowth. mTOR functions in two structurally and functionally distinct complexes, mTORC1 and mTORC2 [19, 30, 31]. Activation of mTORC2 in the brain has not been widely studied. It has been reported that mTORC2, but not mTORC2, could have an important role in the maintenance of the actin cytoskeleton and neurite differentiation [19, 24]. Considering these findings, we think that ATRA may have a role in modulating neurite outgrowth through mTORC2.

In summary, this study demonstrates that traditional synthetic retinobenzoic acid AM580 and novel synthetic retinobenzoic acid RH2-41-2 share the same mechanism of inducing neurite outgrowth in SK-N-SH-N cells. They stimulate neurite outgrowth via activation of both ERK1/2, which results in downstream regulation of RARβ, and mTOR. In contrast to these retinobenzoic acids, ATRA-induced neurite outgrowth is mediated through the activation of ERK, but not mTOR, signaling (Figure 9). Furthermore, according to the structure-activity relationship analysis, differences in hydrophobic group size, tertiary structure and the binding position of functional group result in novel synthetic retinobenzoic acids that show various effects that are mediated by different mechanisms.

**Figure 9.**
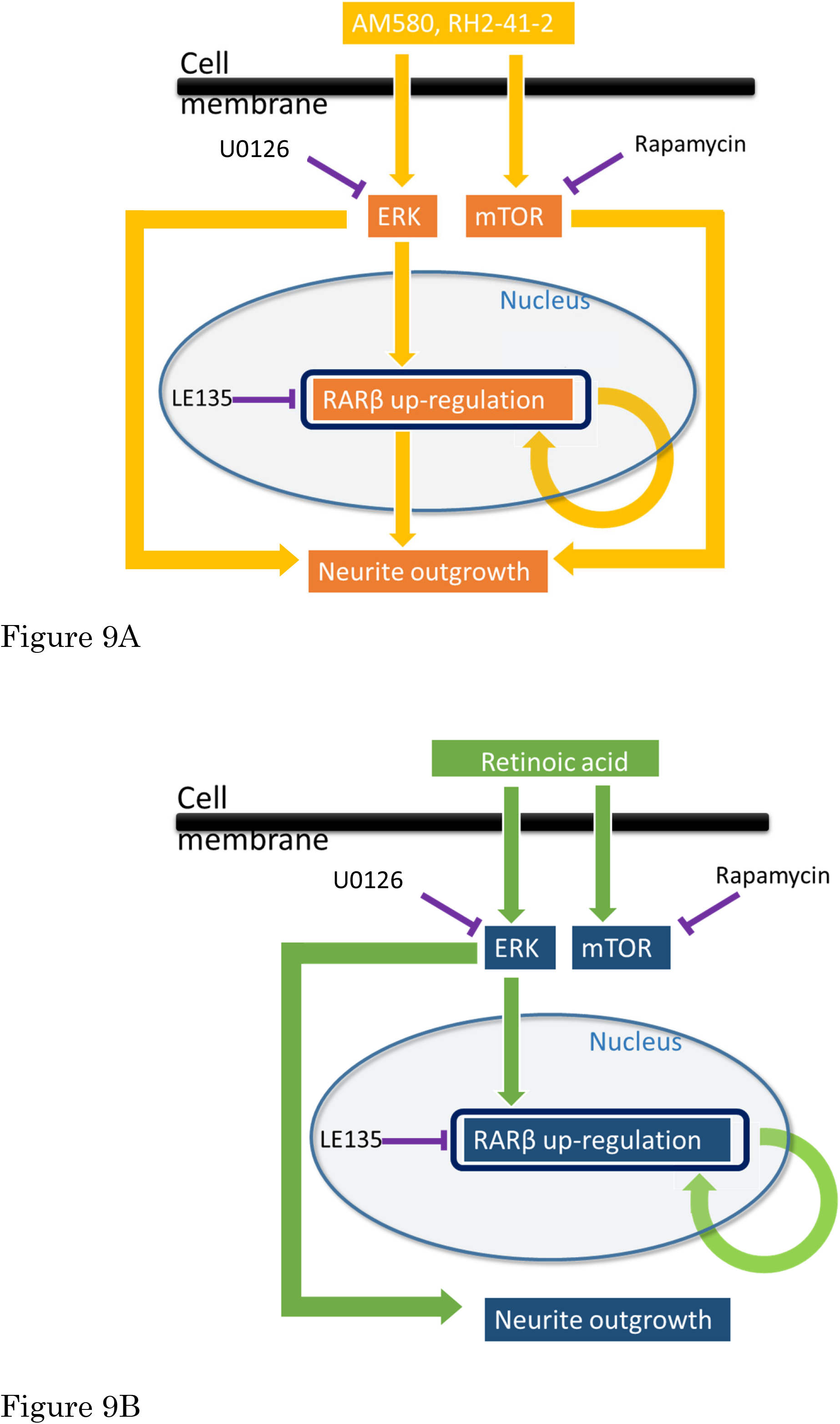
Schematic representation of proposed molecular events involved in neurite outgrowth induction by (A) AM580 and RH2-41-2, or (B) retinoic acid. Arrows mean routes of signaling. “⊣” represents the relationship between the inhibitors and the target.

Finally, it will be important to determine if similar pathways are involved in retinobenzoic acids-mediated differentiation *in vivo*. In addition, it is also important to identify whether these compounds show a lower than RA lipid solubility *in vivo*.

## Acknowledgments

We are grateful to Professor Takahisa Shinomiya (Department of Pharmacy, Faculty of Pharmaceutical Sciences, Aomori University) for his technical support and invaluable discussions.

